# Mutation of *mltG* increases peptidoglycan fragment release, cell size, and antibiotic susceptibility in *Neisseria gonorrhoeae*

**DOI:** 10.1101/2023.08.23.554517

**Authors:** Tiffany N. Harris-Jones, Krizia M. Pérez Medina, Kathleen T. Hackett, Melanie A. Schave, Ryan E. Schaub, Joseph P. Dillard

## Abstract

Infection with the Gram-negative species *Neisseria gonorrhoeae* leads to inflammation that is responsible for the disease symptoms of gonococcal urethritis, cervicitis, and pelvic inflammatory disease. During growth these bacteria release significant amounts of peptidoglycan (PG) fragments which elicit inflammatory responses in the human host. To better understand the mechanisms involved in PG synthesis and breakdown in *N. gonorrhoeae*, we characterized the effects of mutation of *mltG*. MltG has been identified in other bacterial species as a terminase that stops PG strand growth by cleaving the growing glycan. Mutation of *mltG* in *N. gonorrhoeae* did not affect bacterial growth rate but resulted in increased PG turnover, more cells of large size, decreased autolysis under non-growth conditions, and increased sensitivity to antibiotics that affect PG crosslinking. An *mltG* mutant released greatly increased amounts of PG monomers, PG dimers, and larger oligomers. In the *mltG* background, mutation of either *ltgA* or *ltgD*, encoding the lytic transglycosylases responsible for PG monomer liberation, resulted in wild-type levels of PG monomer release. Bacterial two-hybrid assays identified positive interactions of MltG with synthetic penicillin-binding proteins PBP1 and PBP2 and the PG-degrading endopeptidase PBP4 (PbpG). These data are consistent with MltG acting as a terminase in *N. gonorrhoeae* and suggest that absence of MltG activity results in excessive PG growth and extra PG in the sacculus that must be degraded by lytic transglycosylases including LtgA and LtgD. Furthermore, absence of MltG causes a cell wall defect that is manifested as large cell size and antibiotic sensitivity.

**Importance:** *Neisseria gonorrhoeae* is unusual in that the bacteria release larger amounts of cell wall material as they grow as compared to related bacteria, and the released cell wall fragments induce inflammation that leads to tissue damage in infected people. The study of MltG revealed the importance of this enzyme for controlling cell wall growth, cell wall fragment production, and bacterial cell size and suggest a role for MltG in a cell wall synthesis and degradation complex. The increased antibiotic sensitivities of an *mltG* mutant suggest that an antimicrobial drug inhibiting MltG would be useful in combination therapy to restore the sensitivity of the bacteria to cell wall targeting antibiotics to which the bacteria are currently resistant.

## Introduction

*Neisseria gonorrhoeae* is a Gram-negative diplococcus that is the causative agent of the sexually transmitted infection gonorrhea. Treatment of this infection has become increasingly challenging due to antibiotic resistance to all previously used antibiotic therapies, highlighting the need for new treatments and new drug targets (1). Currently, ceftriaxone is the only antibiotic therapy that is recommended for treatment of gonorrhea (2). Serious consequences of gonorrhea include infertility, pelvic inflammatory disease, ectopic pregnancy, neonatal blindness, and disseminated gonococcal infection (3). Symptoms and pathology of the infection are derived from the large inflammatory response that occurs in most gonococcal infections (3). Bacterial products released by gonococci, including lipooligosaccharide, outer membrane vesicles, heptose-containing metabolites, and peptidoglycan, contribute to this inflammatory response (3). During growth, gonococci release significant amounts of peptidoglycan (PG) fragments that are known inflammatory products (4). These PG fragments are sufficient to cause the death of ciliated cells in human Fallopian tube tissue, recapitulating the damage that occurs during gonococcal pelvic inflammatory disease (5–7).

PG consists of repeating subunits of *N*-acetylmuramic acid (MurNAc) and *N-* acetylglucosamine (GlcNAc). A short peptide chain is attached to MurNAc which serves to crosslink adjacent strands of PG. Several enzymes are involved in the breakdown of PG in the cell wall, including lytic transglycosylases, carboxypeptidases and endopeptidases, and an *N*-acetylmuramyl-L-alanine amidase. The combined action of these enzymes results in the production of small PG fragments. Lytic transglycosylases cleave the MurNAc-β-(1,4)-GlcNAc linkage in PG and generate PG monomers, the most abundant PG fragments released by gonococci (8). *N. gonorrhoeae* releases PG monomers in the form of 1,6-anhydro disaccharide tetrapeptide (GlcNAc-anhydroMurNAc-Ala-iGlu-Dap-Ala) and 1,6-anhydro disaccharide tripeptide (GlcNAc-anhydroMurNAc-Ala-iGlu-Dap) (9). Seven lytic transglycosylases have thus far been identified in gonococci (10, 11). Some lytic transglycosylases have specialized roles in the cell, such as LtgC, which is involved in cell separation (12), while others have significant effects on PG fragment release, such as LtgA and LtgD, where monomer release is abolished upon loss of both of these proteins (8).

MltG is a lytic transglycosylase that was previously characterized in *Escherichia coli, Pseudomonas aeruginosa, Vibrio cholerae, Bacillus subtilis,* and *Streptococcus* species (13–19). In *E. coli,* MltG was suggested to be a terminase, stopping the elongation of glycan strands (14). In *B. subtilis* and *E. coli*, MltG was found to interact with both classes of penicillin binding proteins (PBPs) which are involved in PG synthesis (16, 17). Studies in *P. aeruginosa* show an *mltG* deletion can increase susceptibility to antibiotics (13, 20). The role of MltG has not been characterized in gonococci; however, we hypothesize that MltG plays a role in PG biosynthesis similar to that seen in *E. coli* and *B. subtilis* (14, 17). We characterized the role of MltG in PG fragment release, protein-protein interactions, antibiotic sensitivity, PG turnover, and autolysis.

## Results

### MltG enzymatic function

To examine the function of gonococcal MltG, the enzyme was expressed in *E. coli* as a fusion protein with an N-terminal His tag and a C-terminal maltose binding protein fusion. The enzyme was added to radiolabeled gonococcal sacculi, and the products of the reaction were analyzed by size-exclusion chromatography. The sacculi were purified from a gonococcal strain lacking PG acetylation, to allow cleavage by lytic transglycosylases. Chromatographic separation of the soluble PG fragments resulting from MltG digestion of gonococcal sacculi showed that MltG generated large PG fragments, PG dimers, and PG monomers (Fig. 1A). To determine if MltG was producing glycosidically-linked dimers, the MltG-generated PG dimers were digested with lytic transglycosylase LtgD and analyzed by reversed-phase HPLC. The digested products consisted of 1,6-anhydro disaccharide tripeptide and 1,6-anhydro disaccharide tetrapeptide indicating that they were derived from glycosidically-linked PG dimers (Fig. 1B). The production of PG monomers and glycosidically-linked PG dimers was previously observed for sacculus digestion by the related protein in *E. coli* (14). These data are consistent with gonococcal MltG acting as an endo-lytic transglycosylase.

**Figure 1.**
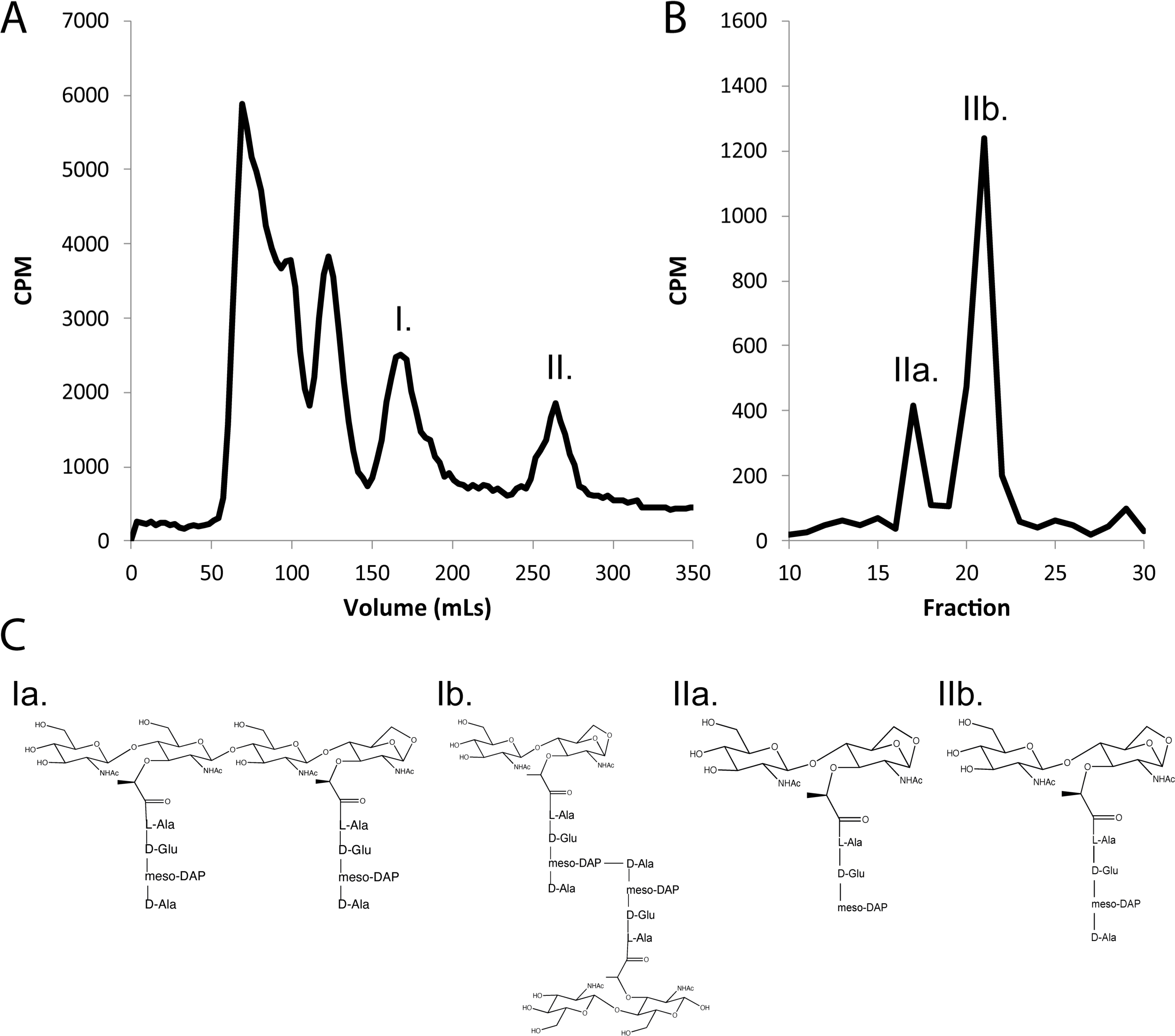
Digestion products of MltG. a) A soluble form of MltG was purified and used to digest whole sacculi metabolically labeled with [6-^3^H]-glucosamine. The soluble PG fragments generated from the digestion by MltG were separated by size-exclusion chromatography. MltG generated large PG fragments, PG dimers (I.), and PG monomers (II.). b) Dimers from the digested sacculi were further analyzed by HPLC after digestion with LtgD. The peaks represent tri- (17 min) and tetrapeptide (21 min) PG monomers which can only be produced by LtgD if the PG dimers are glycosidically linked.

### Cultures of an *mltG* deletion mutant show more large cells

To understand the role of MltG in *N. gonorrhoeae*, we made an in-frame deletion of the *mltG* coding sequence in the chromosome of strain MS11. A complement was created by placing a wild-type copy of *mltG* on the gonococcal chromosome at an unlinked locus, between *aspC* and *lctP* (23, 31).

Mutations affecting cell wall degradation can affect the size of the bacterial cells. In *N. meningitidis*, a mutant lacking the PG deacetylase Ape1 produced larger bacterial cells (35). As Ape1 activity is necessary to allow lytic transglycosylase function, the *ape1* mutant would be unable to degrade PG strands. MltG is thought to act to terminate PG chain synthesis (14). Thus, *mltG* mutants may be deficient in terminating the biosynthetic transglycosylation reaction and may synthesize more cell wall than the WT. To determine if *mltG* cells were altered in size, we performed transmission electron microscopy on gonococcal cells in thin-section. The *mltG* cells appeared larger than those of the WT or complement (Fig. 2A). To quantify the apparent differences, we measured cell size for over 1000 cells greater than 0.5µ for WT, *mltG*, and complement (Fig. 2B). The number of cells counted were 1004, 1041, and 1136 for WT, mutant, and complement, respectively. The number of mutant cells that were 0.5-0.6 or 0.6-0.7 were lower than those of the WT and complement, but this apparent difference did not rise to the level of significance. However, for the largest category of cells, those 1.0 or larger, the percentage of those cells in the population was much larger for the *mltG* mutant (14.5%) than that of the WT (3.1%) or complement strain (3.5%) populations. These results indicate that *mltG* mutant cells have altered cell wall morphology, making bigger gonococcal cells.

**Figure 2.**
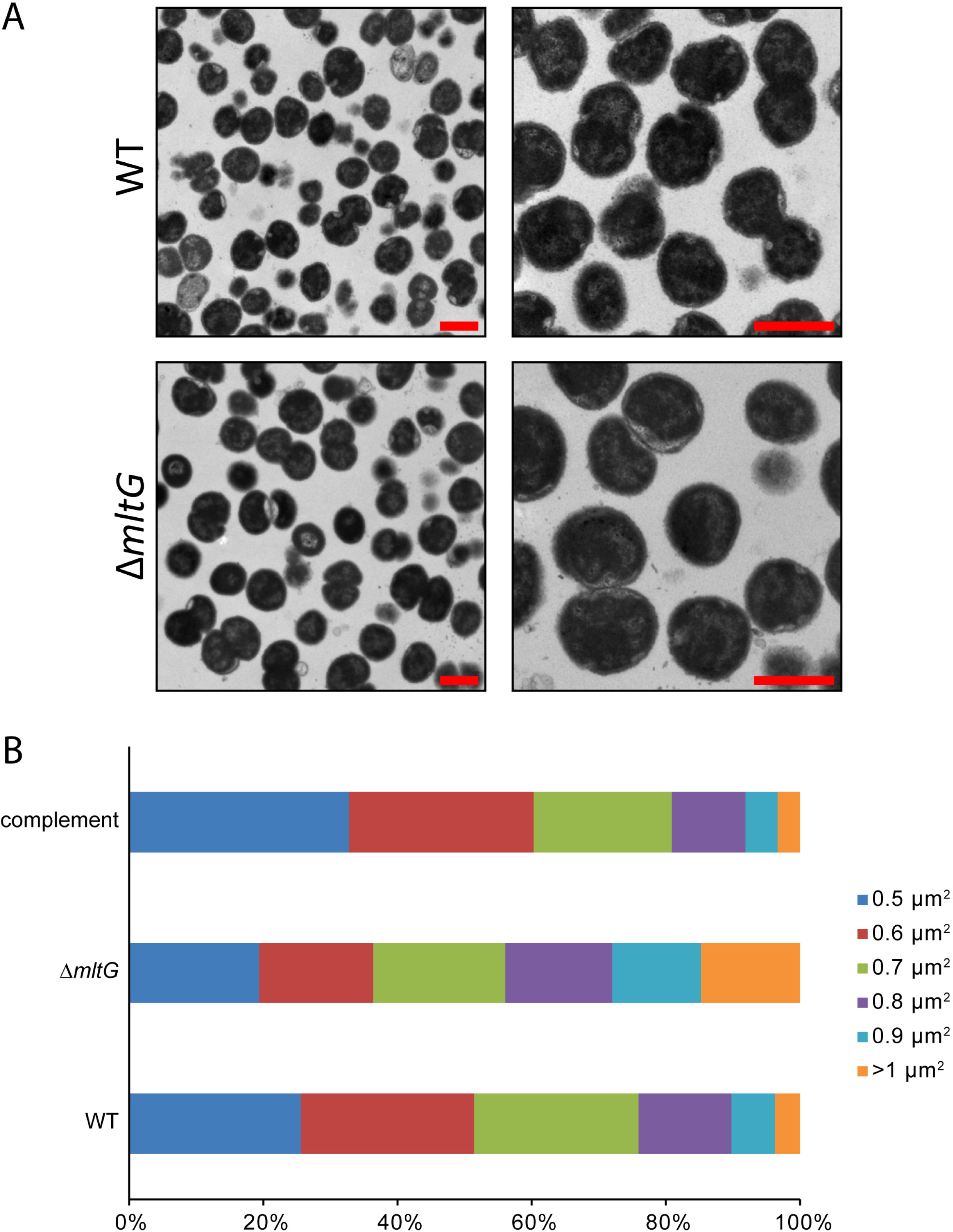
Transmission electron microscopy of *mltG* mutant and WT cells in thin-section. a) Cells of both the WT and deletion (*ΔmltG*) have both mono- and diplococcal cell morphologies. Scale bar = 1 µm. b) The *mltG* deletion has a higher percentage of cells that are larger.

### *mltG* mutation affects PG release and autolysis

The cell wall is an important structure in bacteria for maintaining cell shape and protecting against osmotic stress (38). During growth, PG is constantly being broken down and rebuilt to allow changes in cell size and to allow cell separation (39). A constant balance between degradation and synthesis of the PG occurs to prevent thickening or weakening of the cell wall and subsequent cell lysis. During PG degradation in *N. gonorrhoeae*, most of the PG fragments are recycled back into the cell to be reincorporated into the PG layer (29, 40, 41). We measured PG turnover using metabolic labeling with [6-^3^H]-glucosamine in a pulse-chase experiment. The amount of labeled PG remaining in the sacculus over time was determined. The *mltG* deletion mutant showed a higher rate of turnover, nearly twice that of the wild-type strain (Fig. 3). Complementation restored a wild-type level of PG turnover.

**Figure 3.**
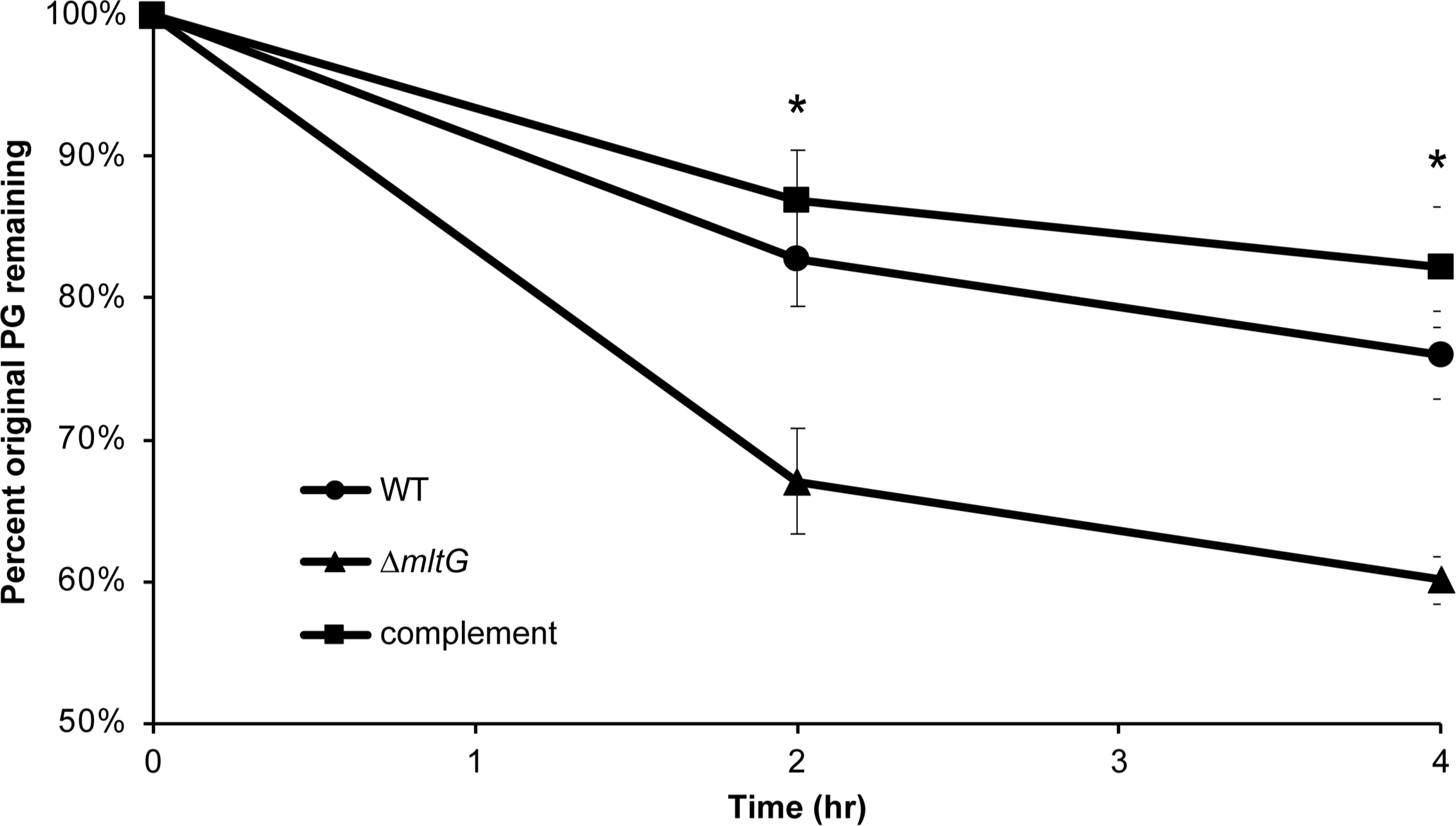
Mutants of *mltG* show a higher rate of PG turnover. *N. gonorrhoeae* strains were labeled with [6,^3^H]-glucosamine and grown in log phase for 4h. At each time-point, cells were removed, the PG in the sacculi was extracted, and radiation measured to determine the amount of original PG remaining in the cell wall. Observations revealed that the *mltG* deletion (*ΔmltG*) had significantly less original PG remaining as compared with WT or complement. * p<0.05

*N. gonorrhoeae* undergoes autolysis when in conditions not favorable to growth (42). To examine the effects of the *mltG* deletion on autolysis, we suspended log-phase gonococci in TrisHCl buffer (pH 8) and measured OD_540_ to follow cell lysis. The *mltG* deletion mutant was significantly less autolytic than the WT in buffer (Fig. 4). Complementation of *mltG* restored the wild-type phenotype. To determine if the decreased lysis in the *mltG* mutant resulted from loss of MltG function or loss of MltG protein, we made a point mutation affecting the predicted MltG catalytic site, *mltG* E213Q. The *mltG* point mutant also showed reduced lysis, although not to the same extent as the deletion mutant. This significant difference in autolysis between the *mltG* mutants and the WT strain might be due to a change in cell wall structure or effects on other peptidoglycanase proteins in the periplasm.

**Figure 4.**
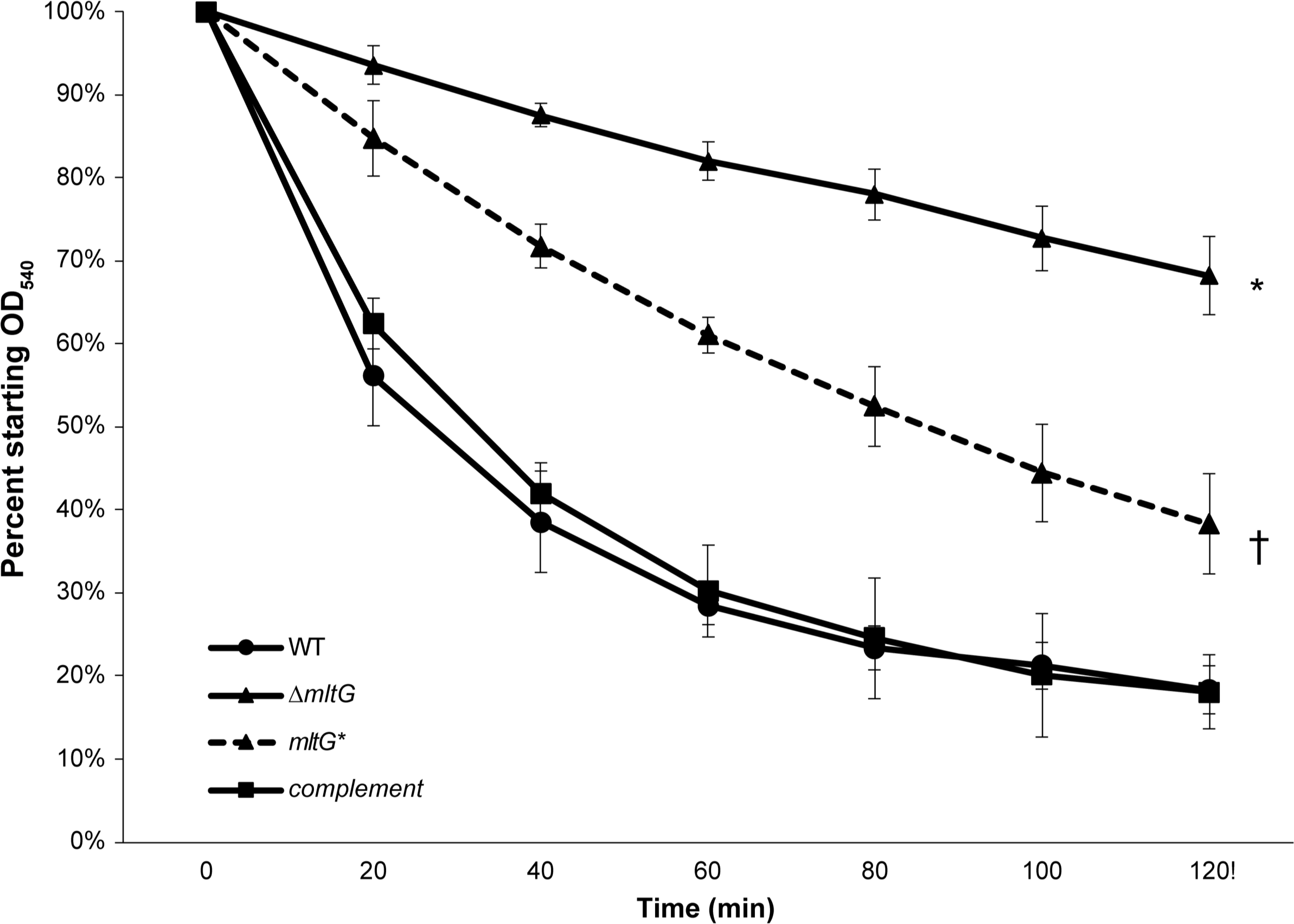
Autolysis in buffer. Strains were grown to log phase and then resuspended in TrisHCl buffer pH 8. Absorbance values at an OD_540_ were taken every 20 minutes for two hours. Both the deletion mutant (*ΔmltG*) (* p<0.01) and point (*mltG\**) mutant († p<0.05) had significantly less lysis compared with wildtype (WT). However, the deletion mutant and point mutant were also significantly different at all timepoints except at 20 minutes (p<0.05). There was no difference in lysis between the wildtype and complement strains.

### Mutants lacking *mltG* release more peptidoglycan monomers and dimers

The PG fragments released by *N. gonorrhoeae* during growth include dimers, monomers, free peptides, free disaccharide, and anhydro-MurNAc (4). Compared with other Gram-negative species, such as *E. coli* or *N. meningitidis*, gonococci release a larger portion of their PG and more of the released fragments are the immunostimulatory monomers and dimers (29, 32). We characterized PG fragment release using pulse-chase metabolic labeling of PG in *N. gonorrhoeae* with [6-^3^H]-glucosamine and separation of the PG fragments released into the supernatant using size-exclusion chromatography (33). The *mltG* deletion mutant was found to release more PG monomer, dimer, and multimer fragments compared to the wild-type strain (Fig. 5A). Complementation restored the mutant to near wild-type levels of PG fragment release. The increased release of PG fragments stands in contrast with phenotypes observed with other lytic transglycosylase defective strains. Mutants lacking *ltgA* or *ltgD* have reduced PG fragment release, and mutants lacking other lytic transglycosylase genes show little to no effect on PG fragment release (8). PG fragment release from the *mltG* point mutant showed a similar phenotype to that of the deletion mutant, with increased amounts of large PG fragments, PG dimers, tetrasaccharide-peptide, and monomers (Fig. 5B).

**Figure 5.**
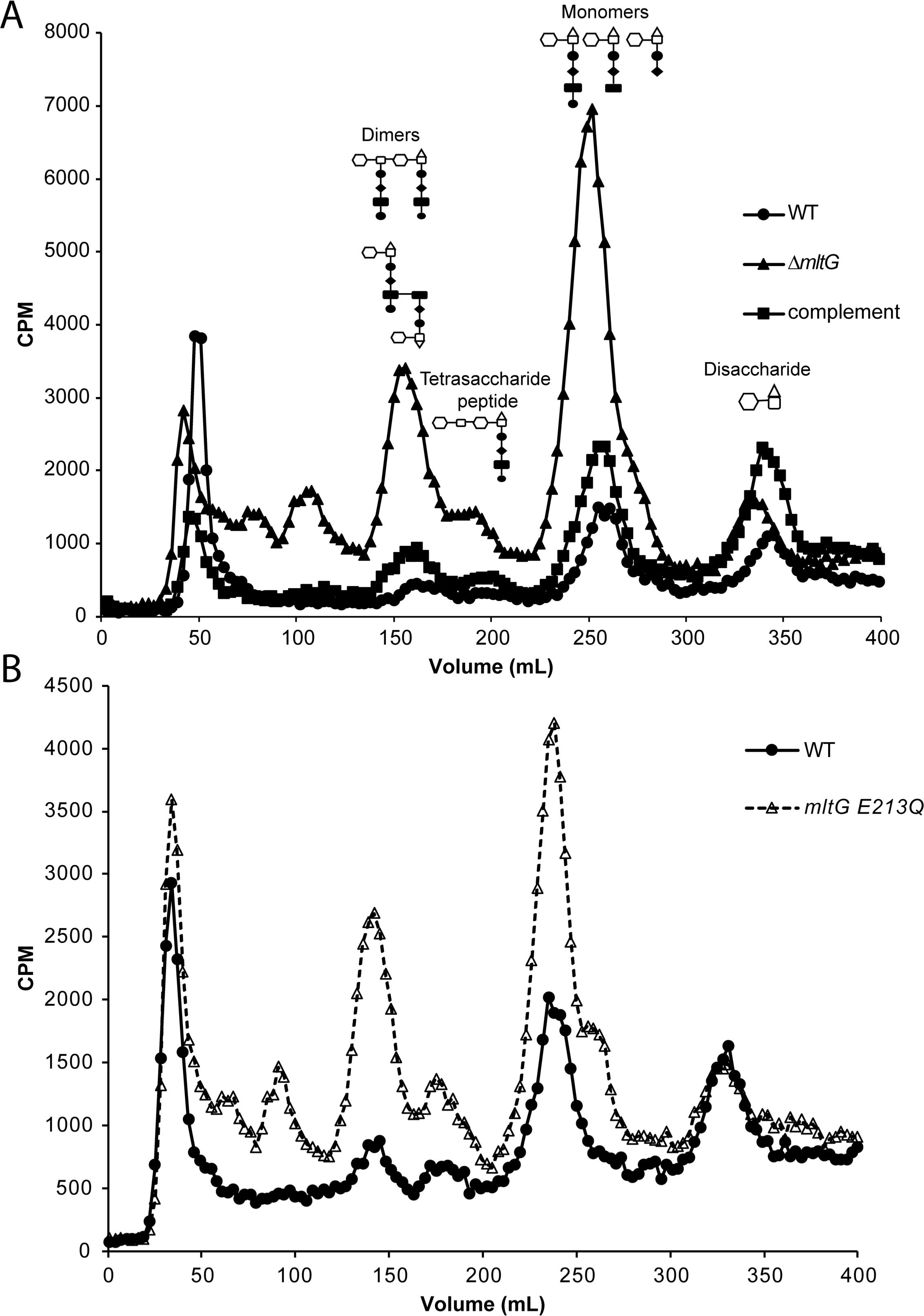
PG fragment release during growth. Cell wall was metabolically-labeled using [6-^3^H]-glucosamine. PG fragments released into the culture supernatant during the chase period were separated by size-exclusion chromatography and quantified as counts per minute (CPM). The strains tested include a strain with an *mltG* deletion (*ΔmltG*), a strain complemented with *mltG* at an ectopic site (complement), and wild type (WT) (A.), and an *mltG* point mutant (*mltG* E213Q) and WT (B.).

### Monomer fragment release is decreased in double mutants lacking *mltG* and other lytic transglycosylases

Both the in-frame deletion and point mutation of *mltG* resulted in a large increase in PG monomer and PG dimer release (Fig. 5). It seems possible that in the *mltG* mutants, excess PG material is being created and is then being degraded by one of the lytic transglycosylases active in creating released PG fragments, i.e., LtgA or LtgD (8, 28). To determine which lytic transglycosylase is responsible for producing the excess of PG fragments released in the *mltG* mutants, we created double mutants with point mutations of the catalytic glutamate for either *ltgA* or *ltgD* plus the point mutation in *mltG.* We found that PG monomer and dimer release was decreased in both the *ltgA mltG* and *ltgD mltG* double mutants compared to the *mltG* single mutant (Fig. 6). Since both double mutants were reduced in amounts of PG monomer fragments released, we cannot assign PG degradation in the *mltG* mutant to just LtgA or LtgD.

**Figure 6.**
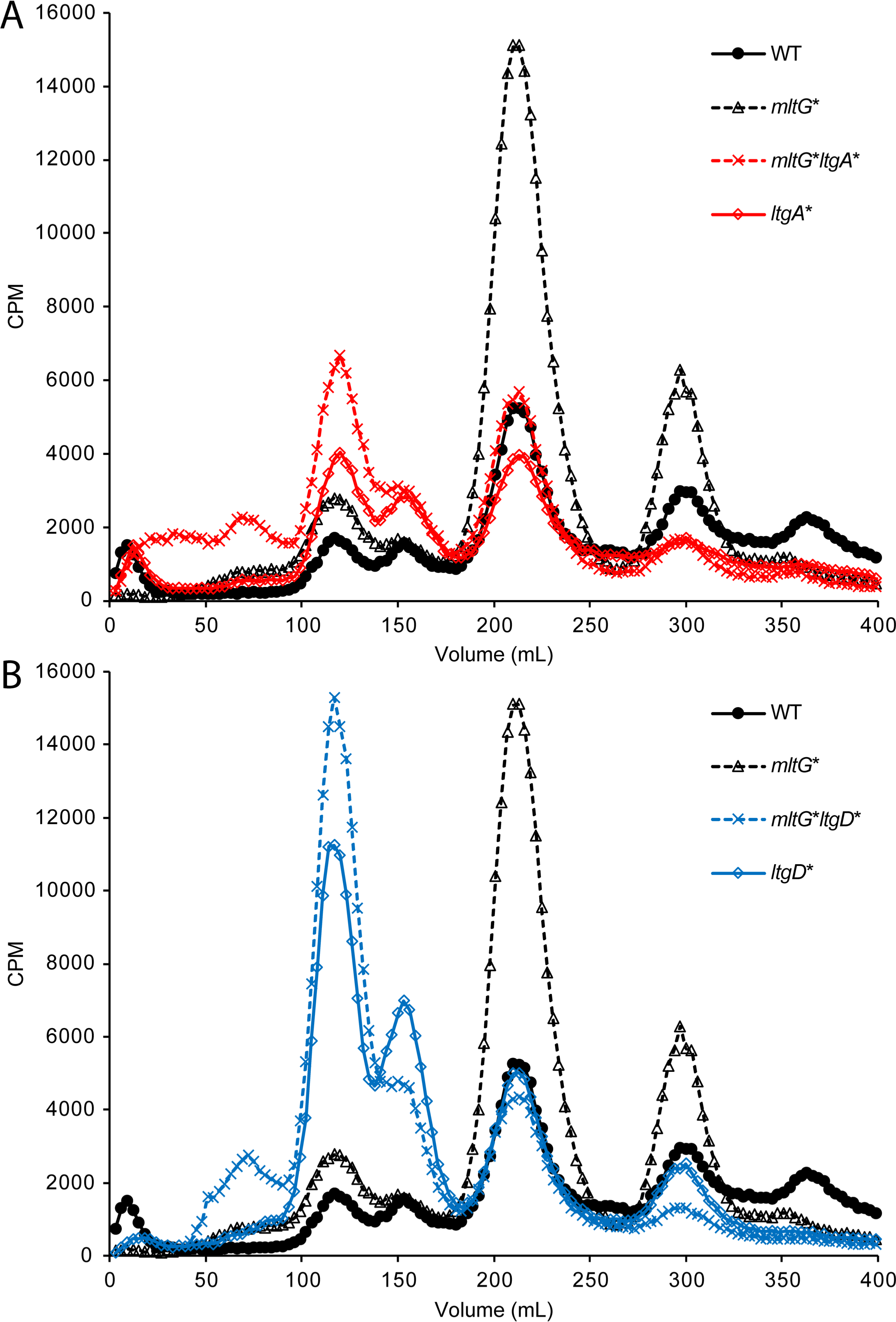
PG fragment release for double mutants of *mltG* and either *ltgA* or *ltgD*. Single and double mutants of *ltgA* (*ltgA\** and *mltG* ltgA\**) (A.) and *ltgD* (*ltgD\** and *mltG* ltgD\**) (B.), were compared to WT and *mltG*\*.

### MltG interactions with PBPs identified by two-hybrid analysis

Synthesis of the cell wall requires the coordination of several proteins to incorporate PG monomers into the existing PG layer (34). In *N. gonorrhoeae*, there are two penicillin-binding proteins (PBPs) involved in synthesis, PBP1 and PBP2, where PBP1 has transglycosylation and transpeptidation activity and PBP2 has only transpeptidation activity (34). PBP1 is responsible for increasing the PG strand length. As PG is assembled in the periplasm, these PBPs bind peptide stems of the newly incorporated PG subunits and crosslink them to the existing cell wall (34). If MltG acts in terminating addition of new PG subunits to the growing PG strand, as has been proposed for other bacterial species (14, 17), then MltG might interact directly with one or both of the biosynthetic PBPs. Using bacterial adenylate cyclase-based two-hybrid assays (BACTH) we identified positive interactions of MltG with both PBP1 and PBP2 (Table 3). MltG preventing continued glycan growth in the cell wall by the biosynthetic complex including PBP1 or PBP2 would stop constant addition of PG monomers to the PG layer. Deletion of *mltG* would be expected to affect PG synthesis by allowing strand synthesis to continue beyond the normal length which can affect cell size, as was seen with some mutant cells (35).

During our BACTH assays we also determined that MltG interacts with PBP4 (Table 3): an endopeptidase and D,D-carboxypeptidase. PBP4 can cleave bonds crosslinking adjacent PG monomers and cleave the terminal alanine residue on the peptide, converting a pentapeptide to a tetrapeptide (36). The other low molecular weight PBP, PBP3, can also perform these functions, and one of these two enzymes must be present for normal separation of PG strands for PG degradation by lytic transglycosylases and AmiC (37). While we obtained positive results for interactions of MltG with PBP1, PBP2, and PBP4, we did not detect interactions of MltG with PBP3, AmiC, LdcA, LtgA, LtgC, LtgD, NlpD, YgaU, YnhG, or YraP.

### MltG mutants are more sensitive to antibiotics that target later processes in PG biosynthesis

Changes in sensitivity to antibiotics can indicate a protein’s importance in a cellular process, and alterations of the cell wall can lead to cell wall-specific antibiotic sensitivities or general defects in permeability. We tested antibiotic resistance to erythromycin, tetracycline, vancomycin, ceftriaxone, fosfomycin, and penicillin G using disk diffusion assays, where the zones of clearing were measured as a representation of growth inhibition. For most antibiotics, the *mltG* deletion mutant had comparable susceptibilities to the wildtype. However, an *mltG* deletion mutant was more susceptible to penicillin, ceftriaxone, and vancomycin, all of which are antibiotics that target cell wall crosslinking (Fig. 7). This result may indicate that an altered cell wall structure in the *mltG* mutant allows more antibiotic to reach its target or that the mutation results in a more permeable outer membrane.

**Figure 7.**
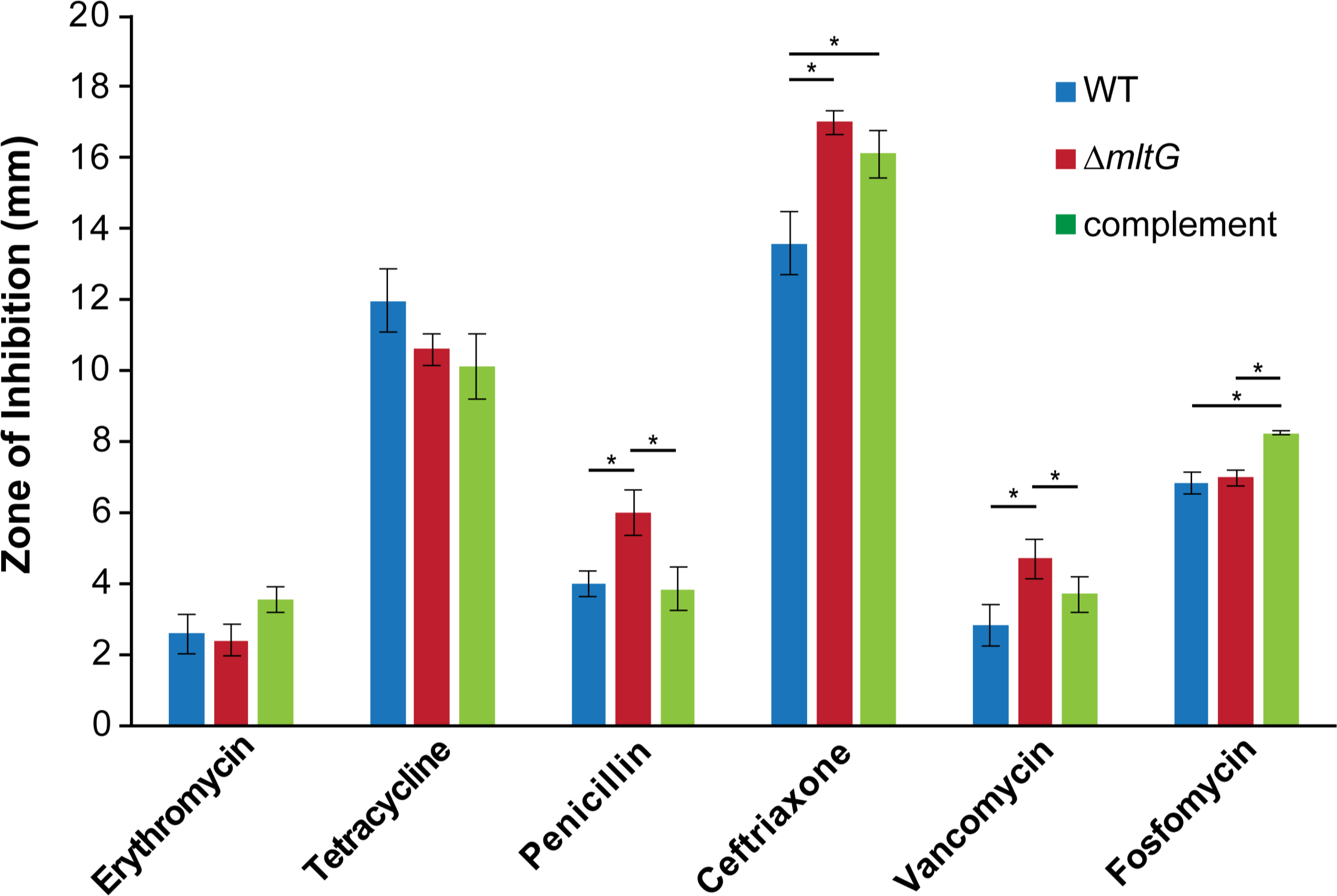
Antibiotic sensitivity. Using disk diffusion assays, the zones of inhibition representing the susceptibility or resistance of a strain to a certain antibiotic were measured. *p<0.05

Although fosfomycin targets cell wall synthesis, there was not a significant change in susceptibility in the *mltG* mutant. The complement strain did show a slight increase in sensitivity, for unknown reasons. Similar to our results, a lack of change in fosfomycin resistance was also observed in a *P. aeruginosa* strain with a deletion of *mltG* (20). Fosfomycin does not target processes involving transpeptidation and PBPs but instead targets a cytoplasmic enzyme. The effects of an *mltG* mutation on antibiotic susceptibility may all be on factors that act at the periplasm or outer membrane.

## Discussion

Lytic transglycosylases play critical roles in several processes in *N. gonorrhoeae* including in PG breakdown, cell separation, and type IV secretion (10–12). MltG is a newly identified lytic transglycosylase in gonococci that has been characterized in other bacterial species. Our goal was to identify the role of MltG in PG metabolism of *N. gonorrhoeae*. When *mltG* is deleted or mutated, more PG fragments are released. This increase indicates that MltG is not involved in monomer production for release. However, the increase in released PG fragments suggests another lytic transglycosylase is involved in producing monomer fragments. To identify this lytic transglycosylase, released fragments from double mutants of *ltgA* and *mltG* and *ltgD* and *mltG* were characterized. Characterizations of released PG fragments for the double mutants showed PG monomer release had been reduced to that of the WT strain, indicating both LtgA and LtgD may be producing the monomers released in the single *mltG* mutant.

Abnormal continued synthesis of PG in the *mltG* mutant may be driving the increased PG turnover. The continued PG synthesis might require increased PG degradation for the bacteria to maintain normal cell size. This hypothesis is supported by the increased number of cells of large cell size. The higher turnover could also indicate a deficiency in recycling, as we have noted previously that certain PG recycling mutants alter their PG fragment uptake in a way that suggests PG fragment monitoring and regulation in *N. gonorrhoeae* (7, 8, 29). Looking further at the cell wall, we also determined that *mltG* mutant cells were less autolytic than the WT under non-growth conditions. The process of autolysis is poorly understood in *N. gonorrhoeae*, but mutants lacking various PG degradation enzymes or lacking a phospholipase are more autolysis-resistant, suggesting that cell wall breakdown or membrane degradation act in this process (43–46). The increased resistance to autolysis in the *mltG* mutant might indicate an altered cell wall structure or decreased autolysis activity of PG-degrading enzymes in the absence of MltG function.

Studying MltG can have importance in combating antibiotic resistance. In *P. aeruginosa* deletion of *mltG* results in decreased MIC or increased sensitivity to ß-lactam antibiotics (20). We similarly found with *N. gonorrhoeae* that an *mltG* deletion changes the sensitivity to this class of antibiotics. The *mltG* mutant was more susceptible to penicillin, vancomycin, and ceftriaxone, which are all antibiotics that target the cell wall and involve the transpeptidation reaction. Targeting MltG could allow the use of antibiotics that were previously off limits due to high levels of resistance. Using this strategy would have large implications on our current struggles to treat gonococcal infection and prevent long term consequences associated with untreatable gonorrhea. It was recently shown that the compound bulgecin A is able to inhibit three different lytic transglycosylases, including MltG, in *P. aeruginosa* (13). Further investigation into this and other lytic transglycosylase inhibitors could be beneficial for future drug development.

The bacterial two-hybrid analysis suggests that MltG interacts with both PBP1 and PBP2. Interactions between MltG and PBPs have also been observed in *E. coli* and *B. subtilis* (14, 17). This result is consistent with a role for MltG in cell wall synthesis. Additionally, MltG was shown to interact with PBP4 which is an endopeptidase in gonococci, similar to PbpG (36, 44). The endopeptidase activities of PBP4 and PBP3 in *N. gonorrhoeae* are critical to normal growth of the bacteria and for function of the amidase AmiC in PG degradation (44, 47). Thus, MltG binding to PBP4 may couple PG synthesis machinery to the PG degradation enzymes that open a space for newly synthesized PG strands.

Through the interaction with PBPs, MltG can terminate elongation through cleavage of PG. The resulting anhydro-muropeptide that caps the strand would prevent further elongation and crosslinking by PBP1 and PBP2. Without this cleavage, PBPs would continue to extend the cell wall growth. As elongation of the glycan chain continues in the absence of MltG, lytic transglycosylases such as LtgA, LtgD, and/or another lytic transglycosylase may attempt to maintain normal cell size by cleaving off excess subunits of PG. As they cleave off excess PG, monomers are released leading to more PG monomer fragment release compared to wildtype. Without MltG, the cell is unable to coordinate synthesis to maintain the normal structure of the PG layer in the cell.

## Materials and Methods

### Bacterial strains and growth

All *N. gonorrhoeae* strains used in this study are derivatives of strain MS11. Piliated strains of MS11 were used for all transformations, whereas non-piliated strains were used for all other experiments. *N. gonorrhoeae* strains were grown at 37°C and 5% CO_2_ on GCB agar plates (Difco) with Kellogg’s supplements (21). Strains were also grown in gonococcal base liquid medium (GCBL) containing 0.042% NaHCO_3_ and Kellogg’s supplements with aeration (22, 23). *E. coli* was grown in lysogeny broth (LB) or on LB agar plates. Antibiotics were used at the following concentrations for *E. coli*: erythromycin at 500 µg/mL, chloramphenicol at 25 µg/mL, and ampicillin at 100 µg/mL. For *N. gonorrhoeae*, chloramphenicol was used at 10 µg/mL, tetracycline was used at 1.5 µg/mL, erythromycin was used at 1.5 µg/mL, ceftriaxone was used at 0.25 µg/mL, fosfomycin was used at 10 µg/mL, penicillin was used at 8 µg/mL, and vancomycin was used at 1.5 µg/mL.

### Plasmid and strain construction

The plasmids used in this study are listed in Table 1. Chromosomal DNA from *Neisseria gonorrhoeae* MS11 was used as a PCR template unless otherwise noted. The primers used in this study are listed in Table 2.

**Table 1.**

| Plasmid or Strain | Description | Reference |
| --- | --- | --- |
| pIDN1/3 | Insertion-duplication plasmid (Erm<sup>R</sup>) | Hamilton et al (2001) (24) |
| pKH37 | Complementation vector (Cm<sup>R</sup>) | Kohler et al (2007) (25) |
| pKH52 | *pacA* point mutation (H329Q) | Dillard and |
|  |  | Hackett (2005) |
| pMRS1 | *mltG* deletion constructed in pIDN3, Gibson cloning | This study |
| pKH189 | *mltG* in pTEV5; 6x HIS tag | This study |
| pKH193 | *mltG* in pKLD116; 6x HIS tag + MBP | This study |
| pKH198 | *mltG* complementation; containing *mltG* in pKH37 (Cm<sup>R</sup>) | This study |
| pKH209 | *mltG* gene block in pIDN3; E213Q point mutant | This study |
| pTEV5 | Vector for synthesis of recombinant protein with a N-terminal hexahistidine, removable by tobacco etch virus (TEV) protease | Rocco et al (2008) (26) |
| pKLD116 | Vector for synthesis of recombinant protein with a N-terminal hexahistidine and maltose binding protein tag in tandem, removable by TEV. | Rocco et al (2008) (26) |
| pRS91 | *ltgA* point mutant constructed in pIDN1 | Schaub et al (2016) (28) |
| pRS92 | *ltgD* point mutant constructed in pIDN1 | Schaub et al (2016) (28) |
| MS11 | Wildtype *Neisseria gonorrhoeae* | Segal et al (1985) (27) |
| MRS500 | MS11 transformed with pMRS1; *mltG* deletion mutant | This study |
| KH530 | MS11 transformed with pKH52; *pacA* point mutation H329Q | Dillard and Hackett (2005) |
| KH624 | KH530 transformed with 1291ΔmsbB chromosomal DNA; *pacA* point mutation H329Q, msbB mutation (Kan<sup>R</sup>) |  |
| KH651 | MRS500 transformed with pKH198; *mltG* complemented (Cm<sup>R</sup>) | This study |
| KH658 | MS11 transformed with pKH209; *mltG* E213Q point mutant | This study |
| KH673 | KH658 transformed with pKH209 and pRS91; *mltG* and *ltgA* double point mutant | This study |
| KH674 | KH658 transformed with pKH209 and pRS92; *mltG* and *ltgD* double point mutant | This study |
| RS555 | MS11 transformed with pRS91; *ltgA* point mutant (E481A) (Erm<sup>R</sup>) | Schaub et al. 2016 (28) |
| RS557 | MS11 transformed with pRS92; *ltgD* point mutant (E158A) (Erm<sup>R</sup>) | Schaub et al. 2016 (28) |
| *Escherichia coli* |  |  |
| Plasmids for BACTH |  |  |
| pKT25 | Cloning vector to add N-terminal T25 fragment for BACTH assays | (Euromedex kit) |
| pUT18C | Cloning vector to add N-terminal T18 fragment for | (Euromedex kit) |
|  | BACTH assays |  |
| pKT25-zip | Positive control for BACTH assays | (Euromedex kit) |
| pUT18C-zip | Negative control for BACTH assays | (Euromedex kit) |

**Table 2.**
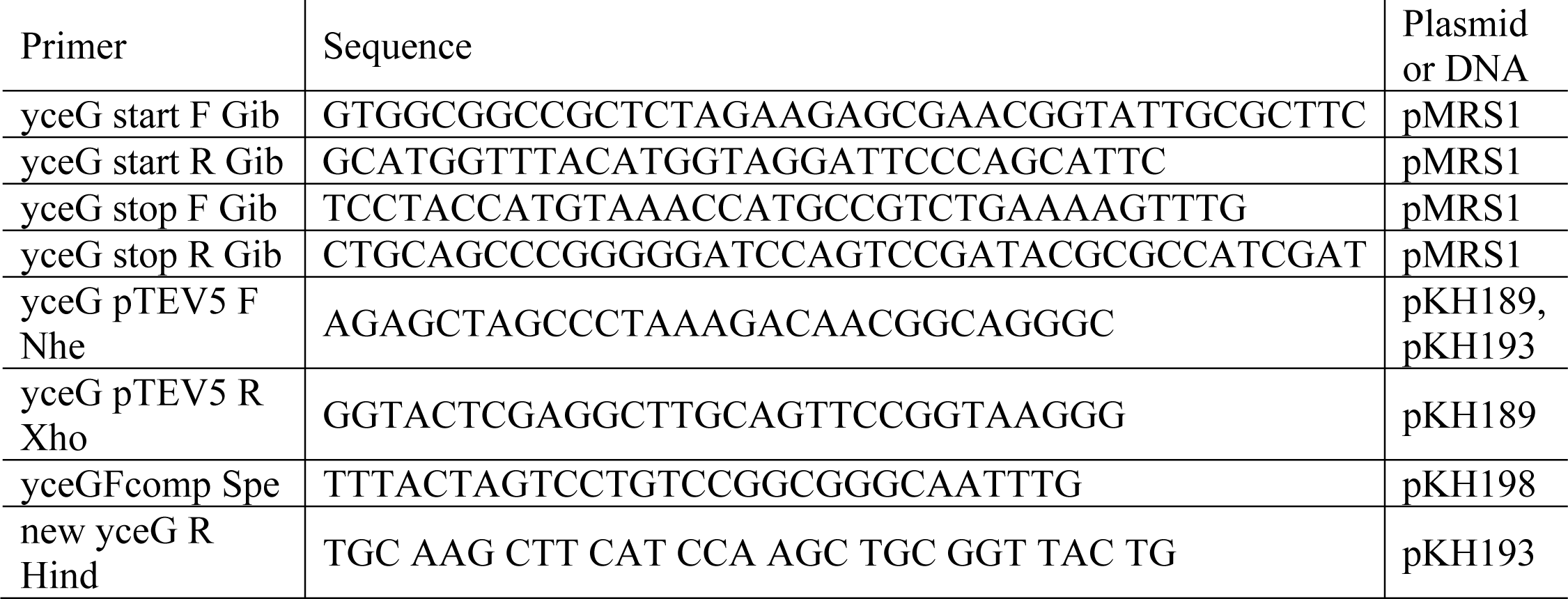

### MltG protein purification

Purification of *mltG* was adapted from Rocco *et al*. (26). An *E. coli* strain expressing pKH193 was induced with 2 mM IPTG at 30°C for 2 hours. Cultures were harvested by centrifugation and washed with column buffer (20 mM NaPO_4_ [pH 7.35], 300 mM NaCl, 20 mM imidazole), and resuspended in 25 mL of column buffer with 0.5% Triton X-100. Resuspended cells were processed with a French press two times to lyse cells. The lysate was then centrifuged for 15 minutes at 20,000 × g. The clarified supernatant was mixed with 250 µL of prewashed nickel resin and mixed by rotation for 1 hour at 4°C. The supernatant-bead mixture was poured into a column and washed with 25 mL column buffer with 20 mM imidazole. The protein was eluted with 500 µL of increasing concentrations of imidazole, 20mM NaPO_4_ pH 7.35, and 300 mM NaCl. Most of protein eluted at 250 mM and 500 mM imidazole. Elution fractions were combined and dialyzed overnight into 50 mM NaPO_4_ pH 7.5, 100 mM NaCl, and 10% glycerol.

### Sacculi labeling and purification

For analysis of PG sacculi, PG from a strain of *N. gonorrhoeae,* with a *pacA* point mutation resulting in a lack of PG acetylation (KH624), was labeled with [6-^3^H]-glucosamine for an hour. The bacterial cells were then harvested by centrifugation and washed once with PBS before resuspension in 500 µL of NaOAc at pH 5. 500 µL of hot 8% SDS was added, and the suspension was boiled for 30 minutes. Sacculi were pelleted by centrifugation, washed with 50 mM sodium phosphate buffer at pH 7.5, and then resuspended in 50 mM sodium phosphate buffer. Isolated sacculi were treated with 100 µg/mL Pronase overnight, to remove any protein from the sacculi. After Pronase treatment, sacculi were boiled in 8% SDS for 30 minutes, then washed twice, before resuspension in water.

### MltG enzyme activity

To test MltG activity, 50 µL of [6-^3^H]-glucosamine labeled, purified sacculi were digested with 25 µL of purified MltG in 25 mM NaPO_4_ buffer pH 6 (total reaction volume was 500 µL) at 37°C overnight. After digestion, MltG was heat inactivated by placing the reaction in a boiling water bath for 10 minutes. To remove insoluble PG, the mixture was centrifuged for 10 minutes at 13,000 rpm. The PG fragments in the sample were subsequently separated using a size-exclusion column. Monomer and dimer PG fragments from the column were collected, concentrated by speed vac, and then desalted prior to HPLC analysis. Tri- and tetrapeptides were separated using a Grace Prevail C18 reversed phase column (250 mm X 4.6 mm) and run over a 4-13% acetonitrile (with 0.05% TFA) gradient for 30 minutes at a 0.5 mL/minute flow rate.

### Disk diffusion assays for antibiotic resistance

*N. gonorrhoeae* was grown overnight on GCB agar. Gonococcal cells were suspended in GCBL medium to an OD_540_ of 0.2. This culture was spread onto GCB plates and incubated for 15 minutes to dry. Diffusion disks were then placed onto the agar, and antibiotics were pipetted onto the disks. Plates were incubated overnight at 37°C in 5% CO_2_. The length, in millimeters, between the edge of the disk and the point where bacterial growth begins was measured. Antibiotics were used at the following concentrations: erythromycin, tetracycline, and vancomycin at 1.5 mg/mL; penicillin at 8 mg/mL; ceftriaxone at 0.25 mg/mL; and fosfomycin at 10 mg/mL. This experiment was repeated three times in triplicate.

### Thin-section transmission electron microscopy (TEM)

For visualization of bacteria by thin-section electron microscopy, strains were grown on GCB plates overnight and then grown in GCBL medium for 3 hours. Cells were harvested by centrifugation, washed once with PBS, and then resuspended in Karnovsky’s fixative. TEM was performed at the University of Wisconsin Medical School Electron Microscope Facility with the assistance of Ben August.

### Autolysis

To measure autolysis of in buffer, gonococci were grown in GCBL medium for 4 hours from an initial OD_540_ of 0.2. Cultures were centrifuged and cells resuspended in 3 mL of 50 mM TrisHCl buffer pH of 6 to wash pellet. Culture was then added to 3 mL TrisHCl buffer pH of 8 in a conical tube at OD_540_ adjusted to 0.3. Cultures were rotated at room temperature and OD_540_ measurements taken every 20 minutes for 2 hours. This experiment was repeated 3 times.

### PG turnover

To assess peptidoglycan turnover during growth, cultures of GCBL medium at an OD_540_ of 0.2 were grown into log phase (3-4 hours). Cultures were then diluted back to an OD_540_ of 0.2 and labeled with 10 µCi/mL of [6-^3^H]-glucosamine for 30 minutes. After labeling, cells were washed once and resuspended into 4 mL GCBL medium. At each timepoint, 1 mL of culture was removed, and 200 µL of *E. coli* culture were added as a carrier. Cells were then centrifuged at 13,000 rpm for 5 minutes. Pellets were at stored at -20°C until analysis. To purify PG, the cell pellets were thawed, resuspended in 0.5 mL NaOAc at pH5, and added to 0.5 mL of hot 8% SDS. The mixture was placed into a boiling water bath for 30 minutes and centrifuged for 30 minutes at 43,000 × g. Supernatants were extracted, resuspended in water and scintillation fluid, and counted for radiation using a Packard Tri-Carb 2100TR liquid scintillation counter.

### Characterization of PG fragment release

Characterization of released gonococcal PG was conducted as described by Cloud and Dillard (45). For labeling the glycan chain, gonococcal cultures were grown in GCBL medium containing 0.4% pyruvate without glucose with 10 µCi/mL of [6-^3^H]-glucosamine. Cultures were labeled for 30-45 minutes. Cultures were then centrifuged, and pellets resuspended in GCBL medium, allowing PG release for 2.5 hours. Cells were removed by centrifugation, and the supernatant was passed through a 0.22 µM filter. Radioactive counts per minute were normalized by taking 60 µL aliquots from each culture and adjusting the culture volumes. This procedure is performed so there are equivalent amounts of radioactivity in the cells in each culture, allowing quantitative comparisons of PG fragment release (29). Supernatants were passed through a size-exclusion column and eluted with 0.1 M LiCl. Fractions were collected consisting of 500 µL of the fraction mixed into 3 mL of scintillation fluid and counted with a Packard Tri-Carb 2100TR liquid scintillation counter or a Perkin Elmer Tri-Carb 4910TR scintillation counter.

### Bacterial adenylate cyclase two hybrid (BACTH) analysis

Bacterial Adenylate Cyclase Two Hybrid System experiments were performed according to the manufacturer’s instructions (Euromedex). Plasmids were created that expressed the protein of interest fused to T25 or T18 fragments. Each pair of plasmids was transformed into chemically competent BTH101 cells, and the culture was incubated in a rotator at 37°C for 1 hour. Following incubation, 100 µL of the transformation mixture was plated onto LB agar plates with 40 µg/mL of kanamycin and 100 µg/mL of ampicillin. Plates were incubated for 2 days at 30°C. For the colorimetric assay, following the 2-day incubation, 3 mL of LB containing 40 µg/mL of kanamycin, 100 µg/mL of ampicillin, 40 µg/mL of X-Gal, and 0.5 mM of IPTG was used. Cultures were grown overnight at 30°C in a tube rotator. The following day, 2 µL of each culture was spotted on LB agar containing 40 µg/mL of kanamycin, 100 µg/mL of ampicillin, 40 µg/mL of X-Gal, and 0.5 mM of IPTG.

**Table 3.** Bacterial two-hybrid analysis of MltG interacts with PG-related proteins. MltG fusions showed positive reactions with PBP1, PBP2, and PBP4. PBP1 is a transglycosylase and a transpeptidase. PBP2 is just a transpeptidase. PBP4 is endopeptidase and a D,D-carboxypeptidase.

| MltG Interactions<br>Proteins Tested | MltG interaction<br>Y/N? |
| --- | --- |
| High-molecular-weight PBPs |  |
| PBP1 | Y |
| PBP2 | Y |
| Low-molecular-weight PBPs |  |
| PBP3 | N |
| PBP4 | Y |
| Amidase-related |  |
| AmiC | N |
| NlpD | N |
| Lytic Transglycosylases |  |
| LtgA | N |
| LtgC | N |
| LtgD | N |
| Other Proteins |  |
| LdcA | N |
| YnhG | N |
| YgaU | N |
| YraP | N |

## Acknowledgements

This work was supported by grant NIH/NIGMS R25GM083252 (T. Harris) and NIH/NIAID R01AI097157 (J. Dillard).

